# MiniMAP: An automatic pipeline for paediatric lower limb musculoskeletal model generation

**DOI:** 10.64898/2026.10.26.761767

**Authors:** Laura Carman, Timothy Salemink Waldren, Thor Besier, Julie Choisne

**Affiliations:** Auckland Bioengineering Institute, The University of Auckland, New Zealand; Department of Engineering Science & Biomedical Engineering, The University of Auckland, New Zealand

## Abstract

The generation of paediatric musculoskeletal models often requires compromise between data availability, accuracy, and computation time. Statistical shape models allow for rapid generation of bone geometries for use in musculoskeletal modelling. An articulated shape model allows for movement at the joints during optimisation to better fit to experimental data. Here we present a methodology for generating custom paediatric OpenSim models using an articulated shape model of the lower limb generated from CT scans of over 300 children aged 4 to 18 years old. Bone geometries are predicted using demographic factors (age, height, mass, and sex), linear bone measurements (bone widths and lengths), and 12 skin surface landmarks (L/R ASIS, L/R PSIS, medial/lateral epicondyles, and medial/lateral malleoli). To account for skin padding (bone to skin surface distance) a random forest algorithm was developed for skin padding prediction for these 12 landmarks. Accuracy of the model prediction was assessed using a leave-one-out analysis to assess landmark errors, bone surface errors, and clinical bone measurement errors (linear measurements and angular values). Motion capture and MRI data were collected from two paediatric test cases to test the workflow on unseen data. Skin padding values varied widely in this population with average values from 3.4mm at the lateral malleolus to 25.9mm at the PSIS. Skin padding was best predicted using age, height, mass, ASIS width, epicondylar width, and malleolar width with an average RMSE of 4.68mm. The articulated shape model predicted accurate bone geometries from skin surface landmarks with average bone surface RMSEs of 2.64 ± 0.99mm, 1.91 ± 0.59mm, and 1.69 ± 0.56mm in the pelvis, femora, and tibiae/fibulae respectively. In the two unseen test cases, average bone surface errors ranged from 1.64 ± 1.2mm to 3.41 ± 3.1mm. This study demonstrated the accuracy of ‘MiniMAP’, for the rapid generation of custom paediatric musculoskeletal models.

**Author Summary:** Accurate modelling of the paediatric musculoskeletal system is important in applications including gait analysis, surgical planning, and rehabilitation. As large datasets in a paediatric population for any type of data are rare, therefore, compromises have to be made in the way both clinicians and researchers analyse motion capture data in a paediatric population. Often, this compromise will involve either a sacrifice in accuracy or a dramatic increase in computational time. For instance, we can approximate paediatric motion using an adult model. While this is a fast method, we know that there are differences when compared to a gold standard model. Alternatively, we can generate a fully subject specific model from medical imaging data, but this is time consuming and costly. Others prefer to use a mix of these two methods. Our research group has been focused on using statistical shape modelling as an alternative approach that allows for both rapid and accurate generation of paediatric musculoskeletal models. Presented here is the latest iteration of that work where we validate the performance of our open-source tools for generating custom models for clinicians and researchers alike.

## 1 Introduction

Childhood musculoskeletal conditions such as cerebral palsy, developmental hip dysplasia, slipped capital femoral epiphysis, and idiopathic torsional deformities significantly impact lower-limb mobility, influencing long-term bone and joint health [1,2]. Personalized musculoskeletal modelling provides a powerful means of examining the interplay between anatomical structure and function, enabling one to link gait data with internal muscle–tendon behaviour and joint loading [3–5]. A key prerequisite for high-fidelity musculoskeletal simulation is accurate anatomical representation including bone geometry, joint alignments, muscle-tendon origins/insertions, and bony landmarks. Definition of these parameters influences joint kinematics and kinetics, and downstream estimates of muscle-tendon lengths, moment arms, and muscle and joint contact forces [5–8].

One common method to generate paediatric musculoskeletal models is to scale an adult template model to subject anthropometry or landmark positions [9]. While this is a fast process, the approach fails to capture paediatric morphology and bony structures, all dependent on growth, making proportional scaling inadequate in a paediatric population [10–13]. Another option for generating paediatric musculoskeletal models is to segment MRI or CT images. Image-based subject-specific models yield anatomical accuracy (considered the gold standard), but are time- and resource-intensive. Manual segmentation and mesh-fitting pipelines for children can take 10–12 h per case and remain impractical for large cohort studies or clinical use [12,14]. Furthermore, most clinical workflows lack the full 3D lower limb scans necessary for generating a model of the complete lower limb. Studies show that generic scaling and image-based models produce different biomechanical outputs in a paediatric population, underscoring the need for a scalable model-personalisation workflow [5,14–16].

Statistical shape models (SSMs) have emerged as a solution to bridge this gap. By capturing population variation via principal component analysis (PCA) and/or partial least squares methods, SSMs can accurately predict individual bone geometries from low-dimensional inputs, such as demographic and length measurements, or motion capture markers in both adult and paediatric populations [10,17–19]. A recent review also underscores the importance of capturing growth-driven morphological variation in paediatric SSMs and selecting clinically accessible predictors [20]. Articulated shape models (ASMs) extend SSMs by modelling multiple connected bones and their relative articulation at the relevant joint, thereby capturing inter-segment geometry and alignment in addition to shape. In adult cohorts ASMs achieved lower prediction errors compared to linear scaling (RMSE = 4.28 mm vs isotropic scaling error ≈ 5.22 mm) using only seven motion capture landmarks [21]. Our group has previously created an ASM to predict the left lower limb from 8 bony landmarks in a paediatric population [22], with RMSEs of 2.63 ± 0.90mm, 1.97 ± 0.61mm, and 1.72 ± 0.51mm for the pelvis, femur, and tibia/fibula, respectively. Linear scaling produced bone surface RMSEs almost twice as high as the ASM, with average RMSEs ranging from 4.38 to 4.79mm. Recent methodological reviews confirm that SSM/ASM approaches are increasingly adopted in orthopaedic and clinical modelling workflows [23], highlighting the importance of developing accurate and clinically relevant tools for both adult and paediatric populations.

Generating a musculoskeletal model from motion capture markers using an ASM requires knowledge of skin padding (the distance between the skin surface and bony landmarks). Variability in skin padding influences the precision of landmark identification to be used for shape model prediction. Studies using ultrasound imaging have demonstrated that skin thickness (including the epidermal, dermal, and subcutaneous layers) varies markedly with age, sex, and anatomical site in the lower limb. Studies of 74 healthy adults and 140 adults with type 1 and 2 diabetes found skin thickness decreased with age, and increased with BMI, with males typically having increased skin thickness compared to females, and differences were found between the left and right lower limb [24,25]. Derraik et al. also included 103 children aged 5 – 19 years old with type 1 diabetes in their dataset and found skin thickness increased with age and BMI. Pubertal onset was also found to strongly affect skin thickness with no sex differences found prior to puberty [24]. Similar results were also found by studies with datasets of 101 children with diabetes [26] and 42 healthy children [27]. In generating regression equations for iliac crest skin padding from 100 healthy young adults aged 16 – 35 years, Town et al. found the main predictors of skin padding to be sex, height, abdomen skinfold, and hip circumference [28]. These findings underscore that both skin padding and thickness is not constant but is modulated by demographic factors and anatomical region. Despite these known variations, no studies have comprehensively mapped skin padding at clinically relevant bony landmarks of the paediatric lower limb. Many modelling workflows assume a constant offset between skin and bone landmarks, yet the evidence suggests that this offset is site- and age-dependent. Quantifying the relationship between skin-surface and bone-surface landmarks therefore becomes a crucial prerequisite for accurate anatomical reconstruction and for interpreting the precision of musculoskeletal models built from skin-based data inputs.

Building on our previous work [10,22] in paediatric SSMs and leveraging evidence of geometry sensitivity and clinical input feasibility, we now extend our approach into four key aims 1) Capture the variation in both sides of the paediatric lower limb using PCA and predict bone geometries from skin surface landmarks using an articulated shape model, 2) determine the relationship between 12 skin surface landmarks (left and right ASIS, PSIS, medial epicondyle, lateral epicondyle, medial malleolus, and lateral malleolus) and bone surface landmarks (skin padding) in a paediatric population aged 4-18 years and predict these values from clinically accessible data, 3) establish the accuracy of the articulated shape model predictions from skin surface landmarks for anatomical landmarks, bone surfaces, clinical bone measurements, and hip joint centres in the lower limb, and 4) automatically generate a musculoskeletal model from articulated shape model outputs using OpenSim’s python API.

## 2 Methods

### 2.1 Bone Segmentation

Post-mortem CT scans of 333 children (137 F, Age: 4-18 years (mean 12 ± 5 years), Height: 96-192cm (mean 148 ± 24cm), Mass: 14-140kg (mean 49 ± 22kg)) were obtained from the Victorian Institute of Forensic Medicine (VIFM, Melbourne, Australia) with ethics approval ‘2023-Choisne-1143-2385/3’ from the VIFM Ethics Committee and ‘AH24671’ from the Auckland Health Research Ethics Committee. This study used retrospective data, which were collected by the VIFM for autopsy purposes between 2006 and 2019. Before autopsy, the VIFM obtained written consent from the individual’s legal guardian. All methods were performed in accordance with the relevant guidelines and regulations. Any participants with ailments affecting the lower limb structure and function were removed from the dataset. The pelvis, femora, and tibiae/fibulae were segmented from the CT scans as in previous work [10,22], to create surface meshes. After removing cases where all the lower limb bones were not visible, the total number of cases used for the articulated shape model was 324 for the left side and 327 for the right side.

### 2.2 Skin surface landmark determination

Bony landmarks were identified for each case in the dataset automatically using the methods outlined in previous work [29]. Skin surfaces were segmented using Mimics Research Suite v27 (Materialise, Leuven, BE) for all available cases in the dataset. The total case number where skin surfaces were segmented and landmarks were calculated was 300; in some cases, the skin surface landmarks could not be determined due to the CT scan being cut off, or the presence of medical equipment in the scan, which blocked the skin surface area where the landmark should be. Skin surface landmarks were determined for the ASIS, PSIS, medial epicondyle, lateral epicondyle, medial malleolus, and lateral malleolus for the left and right sides. These landmarks were found by projecting from the bony landmark along the corresponding axes of the bone as defined in previous work [29] (pelvis: x axis, femur: z axis, tibia/fibula: z axis) until intersection with the skin surface mesh, as shown in Figure 1.

**Figure 1:**
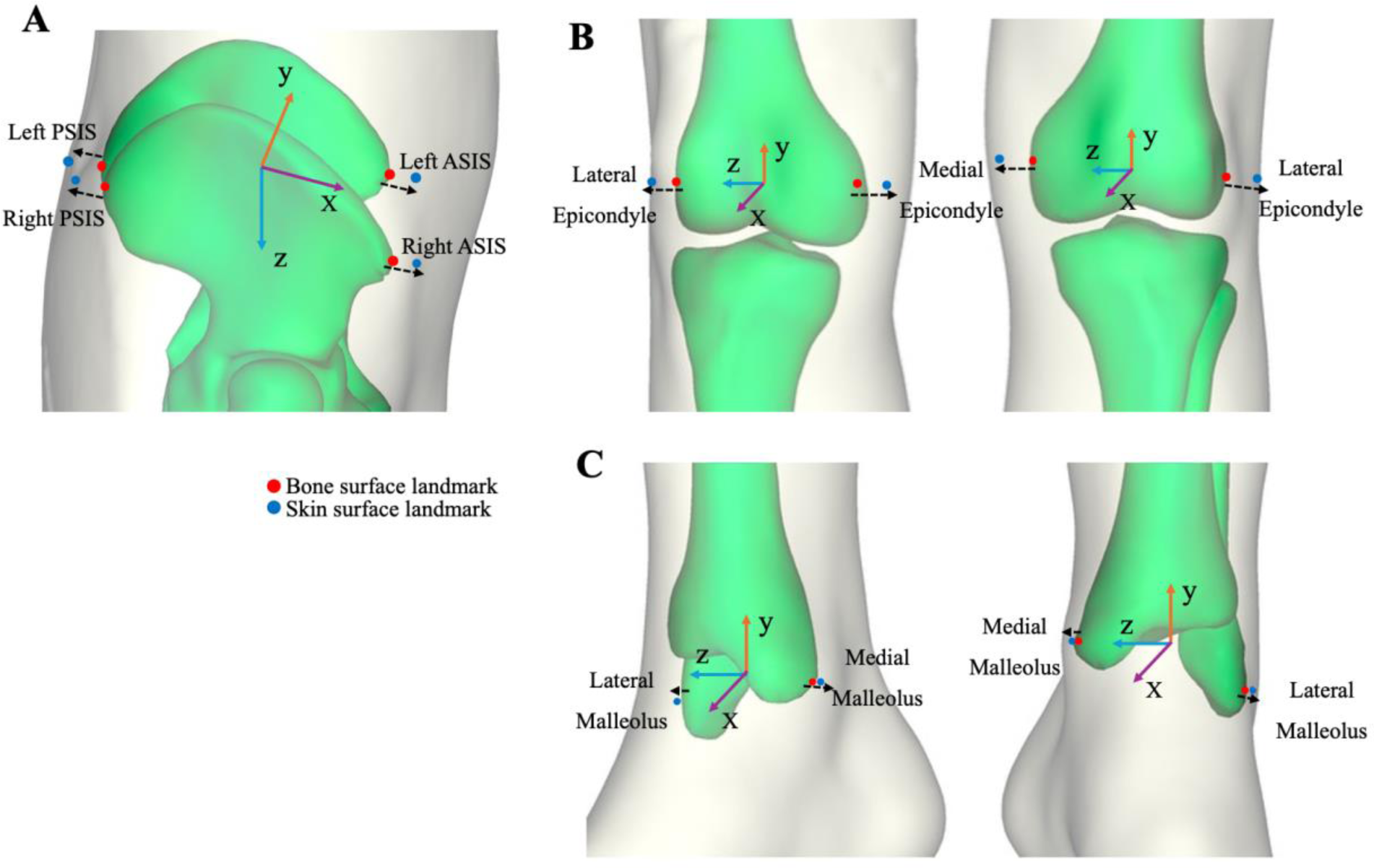
Example case showing skin surface landmarks (blue) determined from bone surface landmarks (Red) for the Pelvis (A), Femur (B), and Tibia/Fibula (C). Landmarks include left/right ASIS and left/right PSIS of the pelvis, medial and lateral epicondyles of the femur, and medial and lateral malleoli of the tibia/fibula.

The relationship between skin surface landmarks and bony landmarks (skin padding) was also determined. Prediction of skin padding was investigated using both Partial Least Squares Regression (PLSR) and Random Forest (RF) algorithms (using sklearn python packages cross_decomposition.PLSRegression and ensemble.RandomForestRegressor). The predictors of skin padding were age, height, mass, sex, ASIS width, epicondylar width, and malleolar width. These predictor variables were chosen as they are easily measured using callipers or derived from motion capture markers in a static, standing reference pose. The best method of prediction and combination of predictors was chosen as the lowest root mean squared error (RMSE) produced using a leave-one-out (LOO) cross-validation.

### 2.3 Articulated Shape model generation

#### 2.3.1 Combined shape model

The articulated shape model for the right side was developed in the same way as the previous articulated shape model generated for the left side lower limb bones in this population [22]. In summary, a combined (or coupled) shape model was developed for the left and the right limb bones separately (as whole pelvis, L/R femur, L/R tibia/fibula). This was performed so the model can account for unilateral differences between sides of the lower limb, for example, in hemiplegic cerebral palsy. The process to generate the combined shape model involves alignment and fitting, using radial basis functions, of the meshes for the pelvis, femur, and tibia/fibula to a template case. These steps achieve nodal correspondence, which is necessary prior to performing PCA. The vertices for the whole pelvis, left femur, and left tibia/fibula were then concatenated into a single mesh file for each case, while retaining the information of which nodes correspond to which bone geometry in a separate file. The same process was performed for the right limb bones. The total number of cases used in the shape model for the left and right side was 324 and 327, respectively. Some cases needed to be removed from the dataset due to missing bone geometries which were unable to be segmented from the CT scans. A PCA was then performed [30] to create the left- and right-side combined shape models. Fitting and PCA were performed using the open-source Python package GIAS3 (https://github.com/musculoskeletal/gias3) [31].

#### 2.3.2 Articulation of the shape model

A detailed description of the articulation of the shape model can be found in our previous publication [22]. Briefly, the articulated shape model predicts lower limb paediatric bones by optimising both shape and pose using the following 12 landmarks: L/R ASIS, L/R PSIS, L/R femoral epicondyles, and L/R tibial malleoli. Landmarks were automatically determined based on previous work [29] and embedded in the left and right shape models as node numbers, with anatomical coordinate systems aligned to be consistent with ISB standards (Figure 2) [32,33]). The pelvis, with origin defined at the midpoint of the ASISs, had six degrees of freedom. Hip joints were defined from a sphere fit to the acetabulum of the pelvis and the femoral head nodes which were aligned to be coincident. The hip joints were modelled as ball-and-socket joints with three rotational degrees of freedom. The knee joint was defined at the centre of the femoral epicondyles and allowed two rotational freedoms (flexion/extension, abduction/adduction), with a 5 mm minimum gap between the femur and tibia to account for cartilage thickness. This addressed varus/valgus alignment and ensured anatomical accuracy. The patella and sacrum were excluded due to limited CT visibility especially in younger children. The articulated shape model predicts bone geometry by minimising the root mean squared distance between the model landmarks and the target landmarks (Figure 3). This was performed using the scipy.optimize.minimize python package with the Powell method to adjust the model PC weights and joint articulation in the following minimisation equation: *x*0 = [*PC_weig_*_ℎ*ts*_, *rigid_rot_*, ℎ*ip_rot_*, *knee_rot_*]. After shape model prediction, bony landmarks and clinical measurements are automatically taken from the predicted meshes following the protocol outlined in [29]. Clinical measurements include torsional values: anteversion angle, tibial torsion, angular measurements: neck shaft angle, femoral and tibial mechanical angles, and linear measurements: width and length measurements from each bone.

**Figure 2:**
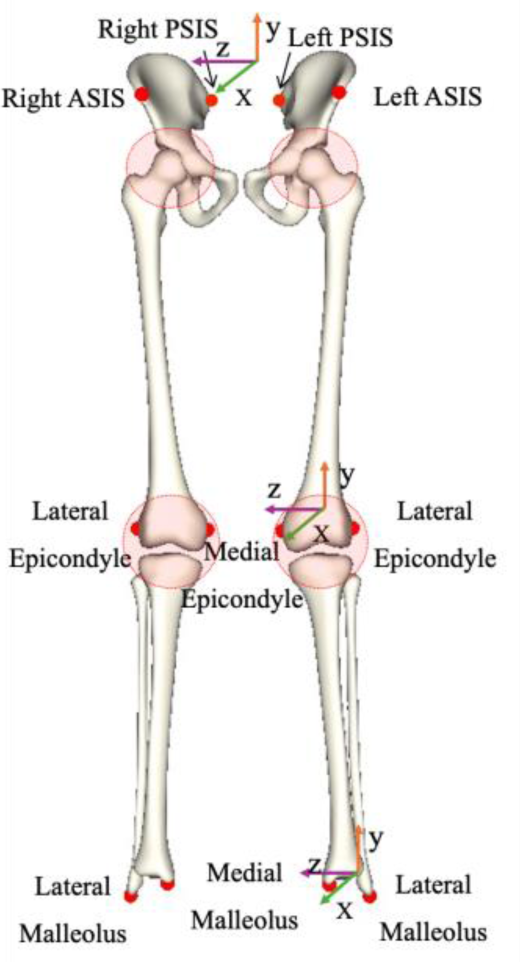
Articulated shape model mean mesh showing coordinate systems and embedded bone surface landmarks for the pelvis (L/R ASIS and PSIS), right and left femur (L/R medial and lateral epicondyles), and the right and left tibia/fibula (L/R medial and lateral malleoli). Left and right hip and knee joints are highlighted in red

**Figure 3:**
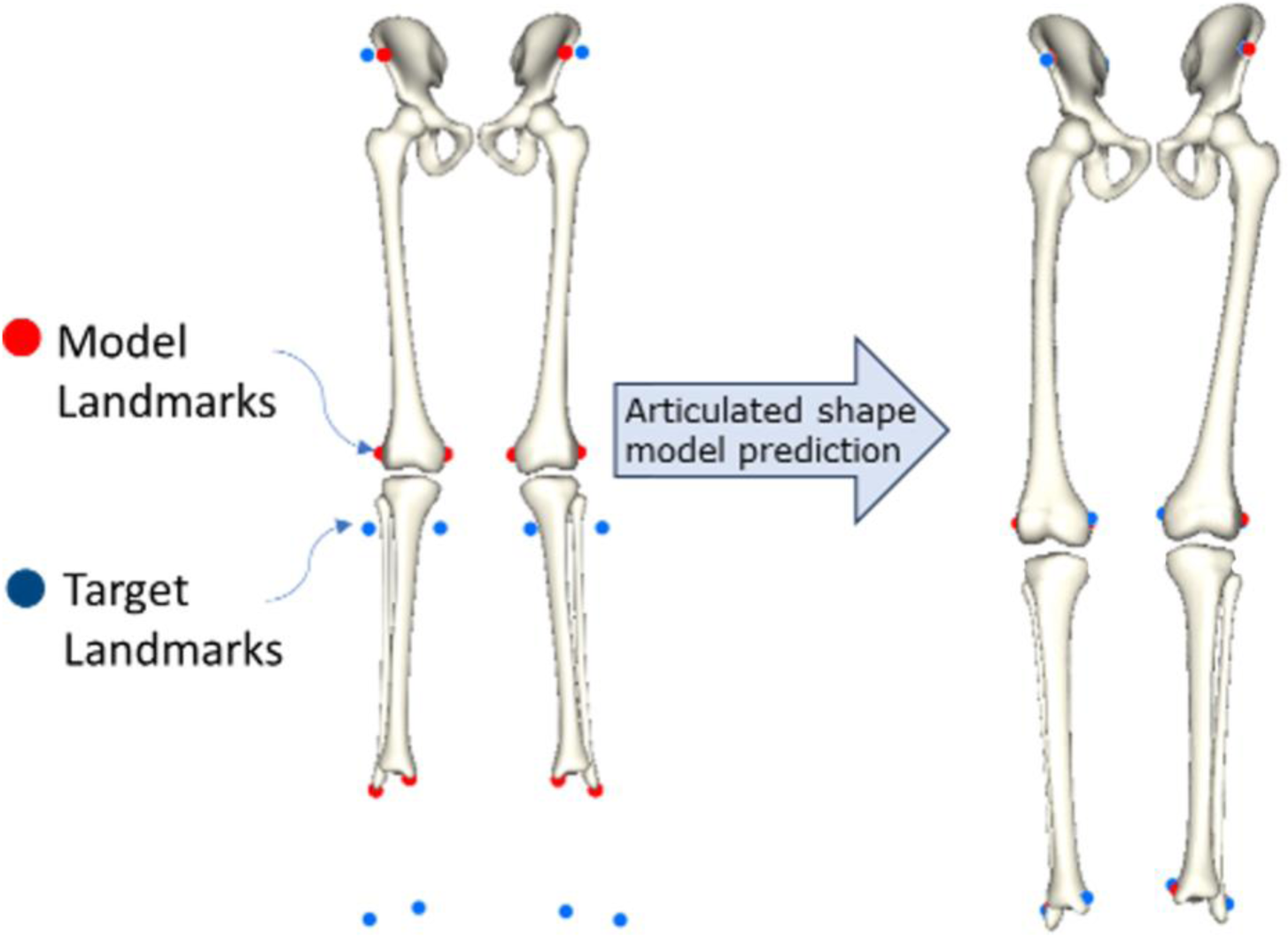
Depiction of how the articulated shape model predicts bone geometry by minimising the distance between the models embedded landmarks (red) and the input target landmarks (blue)

#### 2.3.3 Initial guess and bounds

As an update from our previous model, an initial guess for the PC weights was provided to the shape model using PLSR with the predictors age, height, mass, sex, ASIS width, epicondylar width, and malleolar width. The best combination of predictors was chosen as the lowest root mean squared error (RMSE) produced using a leave-one-out (LOO) cross validation. This was used to bound the PC weights of the model in the minimisation optimisation. Additionally, the joint articulations were bound to be physiologically relevant and not be smaller than those reported in paediatric literature [34–36]. These angles can be seen in Table 1. Overall, these bounds resulted in improved computation time and a lower likelihood of the model producing non-physiological bones, compared to non-bounded optimisation.

**Table 1:** Assigned joint angle bounds for the hip and knee joint of the articulated shape model.

|  | Movement | Degrees |
|---|---|---|
| Hip Joint | Flexion / Extension | 143 / 86 |
|  | Abduction / Adduction | 115 / 45 |
|  | Internal / External Rotation | 92 / 92 |
| Knee Joint | Flexion / Extension | 160 / 12 |
|  | Abduction / Adduction | 9 / 9 |

### 2.4 Validation of ASM predicted bone geometry from skin surface landmarks

To validate the updated articulated shape model, a leave-one-out (LOO) analysis was performed. Where the combined shape model was trained using N-1 participants and predicted the bone geometries of the one ‘left out’ participant using the methodology outlined in 2.3.2 from the skin surface landmarks of each case as determined in 2.2. As the cases included in the left and right shape models were not the same, only participants in which all bone geometry and skin surface meshes were present were included, this totalled 300 cases. The errors were assessed following the same protocol outlined in [22] to determine bone surface errors, bone clinical measurement errors, and bone landmark errors. The performance of the our new articulated shape model (both-limb-ASM) was compared to our previously developed articulated shape model (single-limb-ASM) [22], the key differences being prediction from skin surface landmarks rather than bone surface landmarks, updated model bounding parameters, and prediction of both the left- and right-side lower limb bones. We also compared the new model to our non-coupled individual bone shape statistical shape models of the pelvis, femur, and tibia/fibula (denoted SSM) [10], the key difference being the single bone shape models predicted bone geometry using PLSR from demographic factors and linear bone measurements. The both-limb-ASM was also compared to linear scaling (volumetric based on bone length) of adult bone geometry which was taken from the OpenSim gait 2392 model (denoted LSA) [9]. To assess the difference in errors between the articulated shape model and other prediction methods, a repeated-measure ANOVA was performed.

### 2.5 OpenSim model generation

#### 2.5.1 Coordinate system

A custom OpenSim model [9] was generated using the OpenSim python API. This was built using the predicted bone geometries from the articulated shape model (see 2.3) for the pelvis, femora, and tibiae/fibulae. The model requires the subject’s age, height, mass, and sex as inputs. Firstly, the predicted meshes and marker data were scaled by 0.001 to convert from millimetres to metres. The pelvis was then added to the model as a Body and registered to the OpenSim global coordinate frame by finding the transformation matrix between the pelvis coordinate system (calculated automatically from the pelvis landmarks) and the global frame. The pelvis was attached to the ground using an osim.CustomJoint with the translation set as the pelvis origin (ASIS midpoint) and the orientation set from the calculated transformation matrix (six degrees-of-freedom [dof]). The left and right femur were then added to the model as Bodies with origins at the femoral head centre (calculated as a sphere fit to the femoral head), this point is coincident with the hip joint centre calculated from the pelvis (sphere fit to the acetabulum). The hip joint was modelled using a 3 dof osim.CustomJoint with the flexion axis set as a linear function in the +x direction, the adduction axis set as a linear function in the -z direction (+z for right side), and the internal rotation axis set as a linear function in the -y direction (+y for right side). Additionally, the orientation was set to the joint angles (flexion/extension, ab/adduction, i/e rotation) calculated from the articulated shape model fitting, to effectively ‘zero’ the pose. The left and right tibia/fibula bodies were then added to the model with origins at the midpoint of the malleoli. The knee joint was modelled as a 5 dof osim.CustomJoint with flexion (-x), adduction, and rotation set in the same way as the hip joint. Translation of the knee joint in the x and y axis was controlled using a spline from the OpenSim gait2392 model [9] which models the roll-and-glide of the femur with respect to the tibia/fibula segment. Again, the initial dofs of the knee joint were set to match the joint angles (flexion/extension, ab/adduction) calculated from the articulated shape model fitting. Because we did not have a similar SSM of the foot, we generated feet segments using linear scaling of the Rajagopal Model in OpenSim [37] based on the shank segment length (L/RKNE to L/RHEE). The shank segment length was chosen for foot scaling as researchers and clinicians use various numbers and locations of foot markers for clinical gait analysis. The ankle joint was defined as an osim.PinJoint allowing rotation about the z-axis. The origin of the ankle was set at the midpoint of the malleoli, and the translation and orientation set to match the Rajagopal model.

#### 2.5.2 Default angles

As mentioned in the previous section, default joint angles were initially defined from the articulated shape model fitting process, which involved a registration and fitting of individual bones to best-match the experimental landmark position (including skin padding). These dofs included: hip joint flexion/extension, ab/adduction, and i/e rotation; knee flexion/extension and ab/adduction; and pelvis obliquity, rotation, and tilt.

#### 2.5.3 Body segment parameters (BSPs)

In addition to the body segment morphology and joint descriptions, we also updated the model Body Segment Parameters (BSPs), including each bodies mass, centre of mass location, and the inertial parameters. For children under 14 years old, we followed the regression equations set out by Lahkar et al. [38] to determine the BSPs, derived from a dataset of 668 children, aged 2.9 to 12.7 years old, with balanced sex ratios. For children aged 14 years and older, we used the adult regression equations established by Dumas and Wojtusch [39]. These equations determined segment mass as a percentage of total body mass. The centre of mass location was defined in the coordinate frame of each body. Pelvis length was defined as the average distance between the lumbar joint centre and the centre of the hip joints. The lumbar joint centre was found using the sex-specific regression equations from Dumas et al. [39,40]. The centre of each hip joint was defined using the hip joint centres calculated from a sphere fit to the acetabula of the pelvis. Femoral length was determined from the hip joint centre (calculated as a sphere fit to the femoral head) and the centre of the femoral epicondyles. Tibial length was determined between the midpoint of the femoral epicondyles and the midpoint of the malleoli. The moment of inertia, as radius of gyration in the coordinate frame of each body, was determined as a percentage of segment length. For some BSPs in the under-14 group, the coefficients were also dependent on age.

#### 2.5.4 Knee joint axis optimisation

The knee joint of the model could be further customised using dynamic motion capture data (e.g. walking gait trials). This was performed by updating the opensource code following the methods of Reinbolt et al. [41] from MATLAB (available at https://simtk.org/projects/jmp) to python. This was performed using the scipy.optimize.minimize toolbox while iteratively performing inverse kinematics in the OpenSim API to modify the knee joint orientation about the x-axis (ab/adduction) and the y-axis (i/e rotation). The optimisation aimed to reduce the root mean squared error between model markers and experimental markers by adjusting the flexion-extension axis.

#### 2.5.5 Example cases

To test the accuracy of the articulated shape model and OpenSim model development in a real-world scenario, two participants (Table 2) underwent motion capture testing and a full lower limb MRI scan. Marker trajectories were recorded using a 11-camera motion capture system (VICON Vantage16 cameras, Oxford Metrics, UK). The MRI scan was taken at the Centre for Advanced MRI (CAMRI, The University of Auckland) on a 3T Siemens Skyra scanner (Seimens, Erlangen, Germany) using a sagittal T1-VIBE sequence (T1-weighted fast 3D gradient-echo sequence) with slice thickness: 1.0mm and pixel size: 1.0 x 1.0mm. Bone geometry was segmented using ITK-Snap [42]. Bone geometry was predicted using the articulated shape model from the motion capture landmark locations in the static trial. Landmarks used were left and right ASIS, PSIS, femoral epicondyles, and tibial malleoli. Bone surface geometry and clinical bone measurement errors were compared between ground truth bone geometry (from MRI), prediction of bone geometry using a single bone shape model, and linear scaling.

**Table 2:** Demographics of example cases.

| **Participant** | **Age (yrs)** | **Height (cm)** | **Mass (kg)** | **Sex** |
|---|---|---|---|---|
| **TD02** | 9 | 142 | 29.6 | Male |
| **TD03** | 7 | 133 | 25.8 | Male |

## 3 Results

### 3.1 Bounding of the shape model

#### 3.1.1 Skin padding calculations

A range of skin padding values were found across the subjects for the 12 landmarks except at the ankle (Figure 4). This illustrates the importance of customising skin padding in a predictive model, rather than using an average value. The PSIS markers have the largest range and highest skin padding values, likely due to differences in the orientation of the pelvis (pelvic tilt). Mean skin padding for the pelvis landmarks were 11.0 ± 9.1mm for the ASIS and 25.9 ± 13.4mm for the PSIS, for the femur landmarks were 13.7 ± 4.6mm for the lateral epicondyle and 16.1 ± 6.3mm for the medial epicondyle, and for the tibia/fibula landmarks was 3.4 ± 1.5mm for the lateral malleolus and 4.9 ± 2.2mm for the medial malleolus.

**Figure 4:**
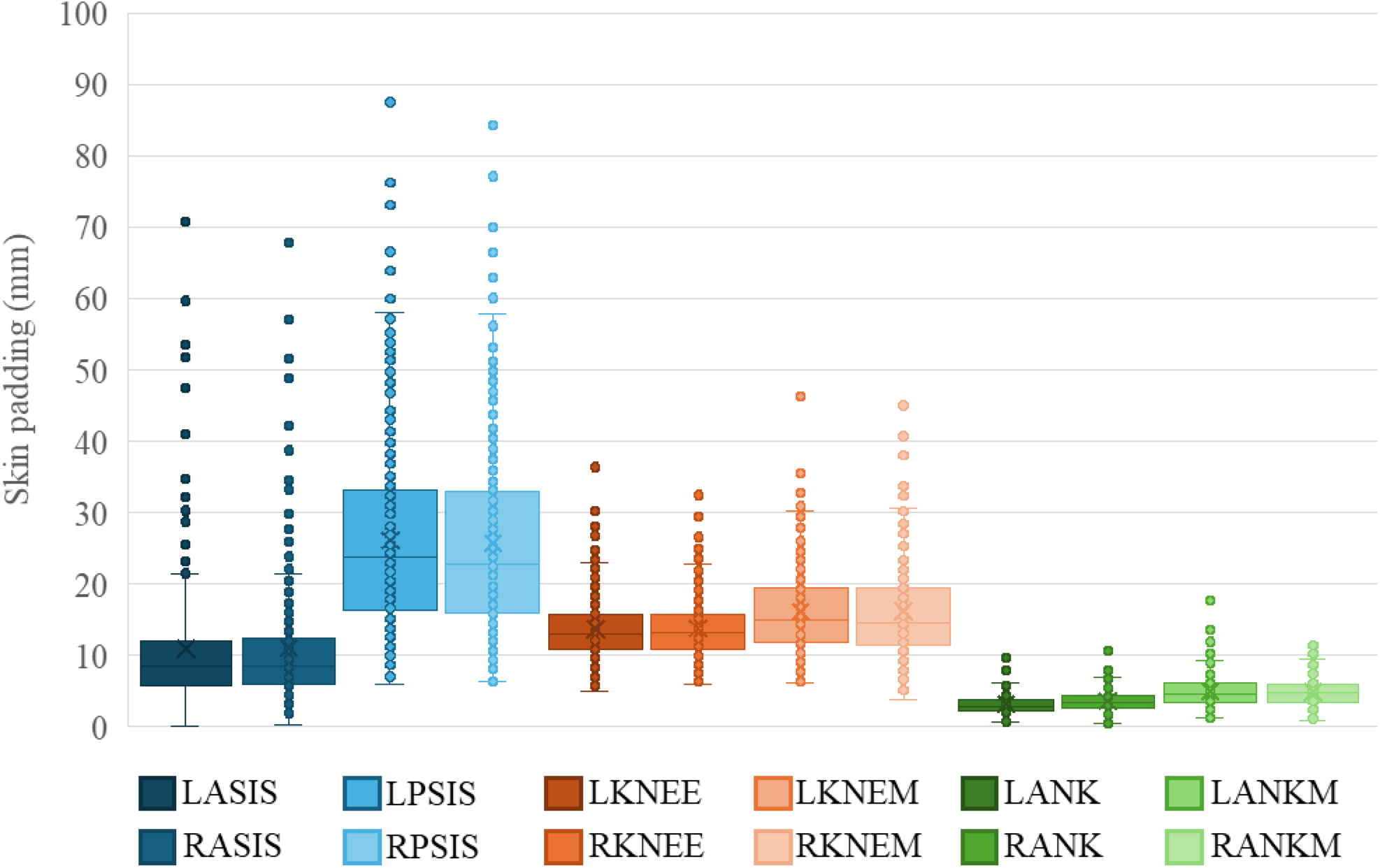
Skin padding (distance between bony and skin surface landmarks) for the L/R ASIS (anterior superior iliac spine), L/R PSIS (posterior superior iliac spine), L/R KNEE (lateral epicondyle), L/R KNEM (medial epicondyle), L/R ANK (lateral malleolus), L/R ANKM (medial malleolus).

The skin padding prediction results for PLSR and RF, using demographic and linear bone measurements as predictors, showed moderate accuracy (Table 3). Overall, the RF algorithm predicted skin padding more accurately for all landmarks and will be used in all future analyses. The RF algorithm produced an R^2^ value of 0.59 and 0.56, and a RMSE of 4.72mm and 4.64mm for the left and right side, respectively. Comparatively, the PLSR produced an R^2^ value of 0.45 and 0.46, and a RMSE of 5.11mm and 5.15mm for the left and right side, respectively. The best predictors for the left side were determined to be age, height, mass, ASIS width, and left epicondylar width. The best predictive factors for the right side were determined to be age, height, mass, ASIS width, right epicondylar width, and right malleolar width.

**Table 3:** Leave one out prediction of skin padding from demographic factors and linear bone measurements using partial least squares regression (PLSR) and random forest (RF) algorithms. Displayed are the best predictive factors, R^2^ of the chosen model, mean absolute error (MAE) ± SD (standard deviation), root mean squared error (RMSE) and normalised root mean squared error (NRMSE)

|  | **Method** | **Predictive Factors** | **R<sup>2</sup>** | **MAE ± SD<br>(mm)** | **RMSE<br>(mm)** | **NRMSE** |
|---|---|---|---|---|---|---|
| **LEFT<br>SIDE** | **PLSR** | *Height, Mass, Sex, ASIS_width,<br>left_epicon_width* | 0.45 | 3.38 ± 3.97 | 5.11 | 5.95% |
|  | **RF** | *Age, Height, Mass, ASIS_width,<br>left_epicon_width* | 0.59 | 2.97 ± 3.68 | 4.72 | 5.40% |
| **RIGHT<br>SIDE** | **PLSR** | *Height, Mass, ASIS_width,<br>right_epicon_width* | 0.46 | 3.36 ± 3.91 | 5.15 | 6.13% |
|  | **RF** | *Age, Height, Mass, ASIS_width,<br>right_epicon_width, right_malleolar_width* | 0.56 | 2.96 ± 3.57 | 4.64 | 5.52% |

#### 3.1.2 Using PLSR to predict an initial guess of the PC weights for the ASM

In constructing the PLSR to predict the PC weights from demographic and linear bone measurements, it was determined that the first PC was predicted well, however the remaining PCs were not predicted as accurately. PC2 produced a smallest NRMSE value of 10.5%, PC3 produced a smallest NRMSE value of 8.5%, and PC4 produced a smallest NRMSE value of 13.1%. Therefore, we decided to only predict the first principal component using PLSR to bound the model and let the minimisation algorithm in the articulated shape model determine the rest of the PC values without bounds. Shown in Table 4 is the prediction of the first principal component using the PLSR algorithm, which produced an R^2^ value of 0.98 and 0.98, and a NRMSE of 3.48% and 3.40% for the left and right side, respectively. The best predictors for both the left and right side were determined to be height and ASIS width.

**Table 4:** leave one out prediction of the first principal component (PC1) from demographic factors and linear bone measurements using partial least squares regression (PLSR). Displayed are the best predictive factors, R^2^ values, and normalised root mean squared error (NRMSE)

|  | **Predictive Factors** | **R<sup>2</sup>** | **NRMSE** |
|---|---|---|---|
| **LEFT SIDE** | *Height, ASIS_width* | 0.98 | 3.48% |
| **RIGHT SIDE** | *Height, ASIS_width* | 0.98 | 3.40% |

### 3.2 Accuracy of the articulated shape model prediction

#### 3.2.1 Bone surface errors

Overall, the articulated shape model accurately predicted the geometry from skin surface landmarks (Table 5). The pelvis showed the highest average RMSE at 2.64mm ± 0.99mm, followed by the femur with 1.91mm ± 0.59mm, and the tibia/fibula at 1.69mm ± 0.55mm. Figure 5 compares the performance of the both-limb articulated shape model (both-limb-ASM) to our previously developed shape models (single-limb-ASM, and SSM) and linear scaling of adult bone (LSA). Our newly developed articulated shape model (both-limb-ASM) performed similarly to the old articulated shape model, indicating the model can just as accurately predict from skin surface landmarks as bone surface landmarks (p = 0.451, 0.225, and 0.715 for the pelvis, femur, and tibia/fibula, respectively). Compared to the single bone shape models (SSM) the updated articulated shape model (both-limb-ASM) produced on average significantly lower bone surface errors (p < 0.001). Compared to linear scaling of adult bone (LSA), the both-limb-ASM produced on average significantly lower bone surface errors (p < 0.001), indicating the importance of more personalised bone geometry in paediatric populations.

**Figure 5:**
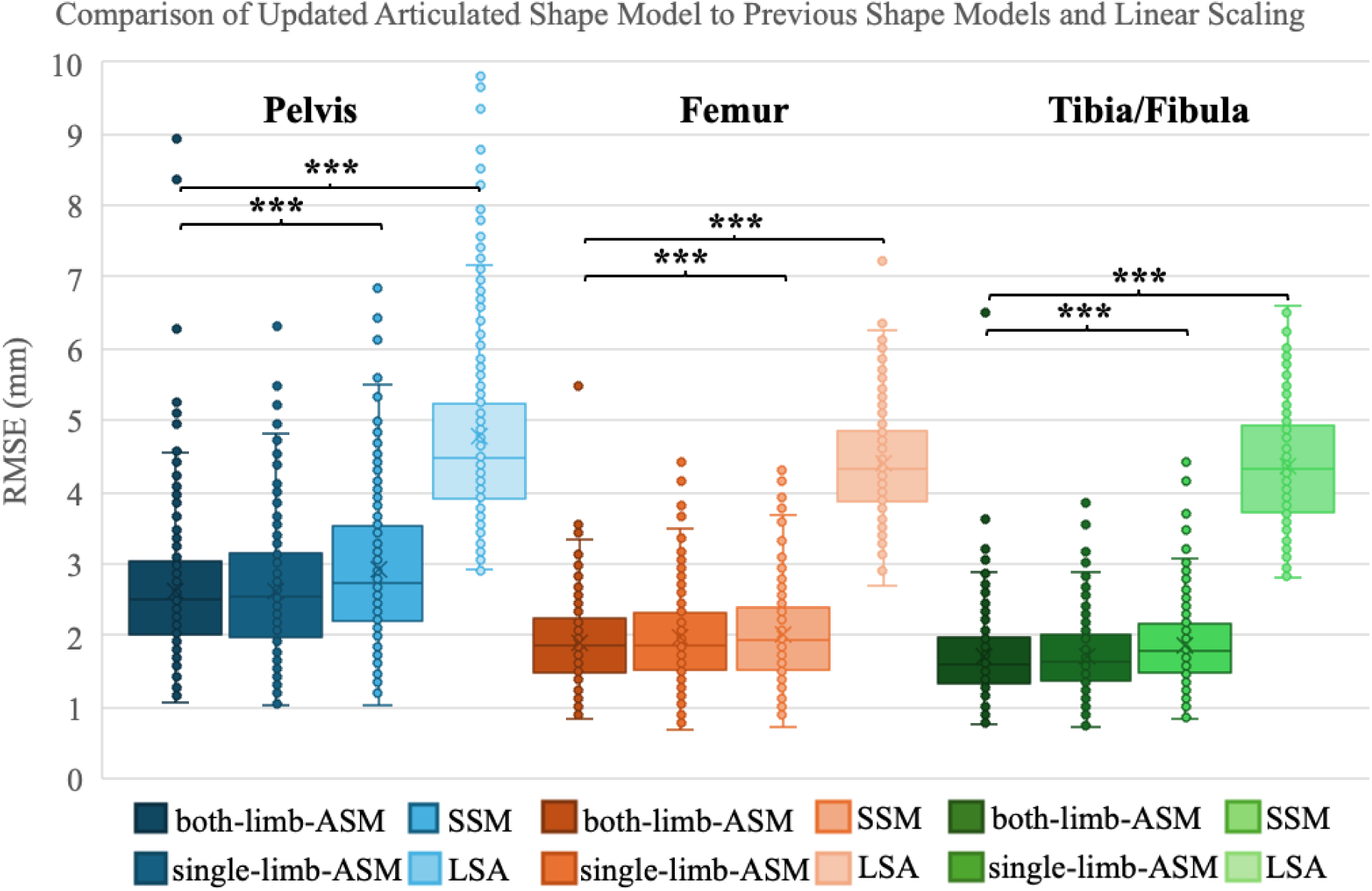
Box plot of bone surface geometry errors for the pelvis, femur, and tibia/fibula predicted from skin surface landmarks using the articulated shape model (both-limb-ASM), compared to errors when predicted from bone surface landmarks using the articulated shape model (single-limb-ASM), errors when predicted using PLSR in single bone shape models (SSM), and errors when performing linear scaling of adult bone geometry (LSA).

**Table 5:** Bone geometry surface errors for the pelvis, femur, and tibia/fibula predicted from skin surface landmarks using both-limb-ASM. Presented as average root mean squared error (RMSE) ± SD, average mean absolute error (MAE) ± SD, and average maximum error (Max) ± SD.

|  | **RMSE (mm)** | **MAE (mm)** | **Max (mm)** |
|---|---|---|---|
| **Pelvis** | 2.64 ± 0.99 | 2.09 ± 0.78 | 9.11 ± 3.26 |
| **Left femur** | 1.90 ± 0.58 | 1.49 ± 0.45 | 6.80 ± 2.10 |
| **Left tibia/fibula** | 1.69 ± 0.56 | 1.32 ± 0.43 | 6.45 ± 2.41 |
| **Right femur** | 1.91 ± 0.59 | 1.49 ± 0.47 | 6.93 ± 2.17 |
| **Right tibia/fibula** | 1.68 ± 0.54 | 1.30 ± 0.42 | 6.67 ± 2.84 |

#### 3.2.2 Bony landmark errors

Bony landmark prediction errors using the both-limb-ASM are shown in Table 6. RMSE in the pelvis landmarks ranged from 8.13 to 8.31mm, in the femur from 4.72 to 4.79mm, and in the tibia/fibula from 5.72 to 7.24mm. The hip joint centre (as a sphere fit to the acetabulum of the pelvis) was predicted with a RMSE of 4.37mm.

**Table 6:** Landmark errors (distance between predicted and ground truth bone surface landmarks) for the ASIS (anterior superior iliac spine), PSIS (posterior superior iliac spine), hip joint centre (pelvis), femoral head centre, greater trochanter, lateral epicondyle, medial epicondyle, posterior lateral condyle, posterior medial condyle, lateral malleolus, and medial malleolus. Errors presented as root mean squared error (RMSE), mean absolute error (MAE), and standard deviation (SD).

| **Landmark** | **RMSE (mm)** | **MAE (mm)** | **SD (mm)** |
|---|---|---|---|
| **ASIS** | 8.31 | 6.76 | 4.84 |
| **PSIS** | 8.13 | 7.02 | 4.10 |
| **Hip Joint Centre** | 4.37 | 3.76 | 2.22 |
| **Lateral Epicondyle** | 4.72 | 4.20 | 2.17 |
| **Medial Epicondyle** | 4.99 | 4.14 | 2.73 |
| **Lateral Malleolus** | 7.24 | 6.31 | 3.56 |
| **Medial Malleolus** | 5.72 | 5.02 | 2.74 |

#### 3.2.3 Clinical bone measurement errors

Clinical bone measurement errors in the both-limb-ASM predictions were similar to the previous iteration of our articulated shape model (single-limb ASM) (Table 7) [22]. These errors were reasonably low for all measurements. Higher errors in linear bone measurements of 5 to 6mm can be seen in PSIS width, femoral length, and tibial length. This is likely due to the range of skin padding values observed at the PSIS landmarks and the lack of information at the proximal femur and tibia, which hinders predictions. The femoral anteversion angle and neck shaft angle prediction errors in the both-limb-ASM were 7.43° and 4.72° on average, respectively, which was also found in the previous iteration of our model (single-limb-ASM). Again, this is likely due to the lack of information at the proximal femur. Notably, errors in tibial torsion and mMPTA decreased by 2.85° and 2.3° on average, respectively, when comparing the both-limb-ASM to the single-limb-ASM.

**Table 7:** Clinical bone measurement mean absolute errors (MAE) for the pelvis, femur, and tibia/fibula predicted using the articulated shape model. Where mLDFA: mechanical Lateral Distal Femoral Angle, and mMPTA: mechanical Medial Proximal Tibial Angle.

| **Measurements** | **MAE (mm)** | **SD (mm)** |
|---|---|---|
| **ASIS width** | 2.13 | 4.28 |
| **PSIS width** | 5.13 | 6.47 |
| **Pelvis Depth** | 3.15 | 4.00 |
| **Femoral Head Diameter** | 1.64 | 2.08 |
| **Epicondylar Width** | 2.64 | 3.49 |
| **Femoral Length** | 5.86 | 7.34 |
| **Tibial Condyle Width** | 2.77 | 3.42 |
| **Malleolar Width** | 2.76 | 3.43 |
| **Tibial Length** | 5.37 | 6.43 |
|  | **MAE (Deg)** | **SD (Deg)** |
| **Anteversion Angle** | 7.43 | 10.25 |
| **Neck Shaft Angle** | 4.72 | 6.34 |
| **mLDFA** | 1.59 | 2.14 |
| **Tibial Torsion** | 4.75 | 6.01 |
| **mMPTA** | 1.30 | 1.76 |

### 3.3 Example cases

The two paediatric test cases for the workflow performed well. Bone surface errors are shown in Table 8, with mean absolute errors in the pelvis ranging from 3.28 to 3.41mm, in the femur from 1.64 to 2.07mm, and in the tibia/fibula from 1.78 to 2.51mm. This is slightly higher than the errors observed in 3.2.1, as expected, since bone geometries were predicted from motion capture landmarks rather than skin surface landmarks. Figure 6 shows that the model had difficulties in predicting the pelvis geometry with regards to pelvis width for both cases. The femur and tibia/fibula predicted geometries closely resemble the ground truth, apart from the distal tibia.

**Figure 6:**
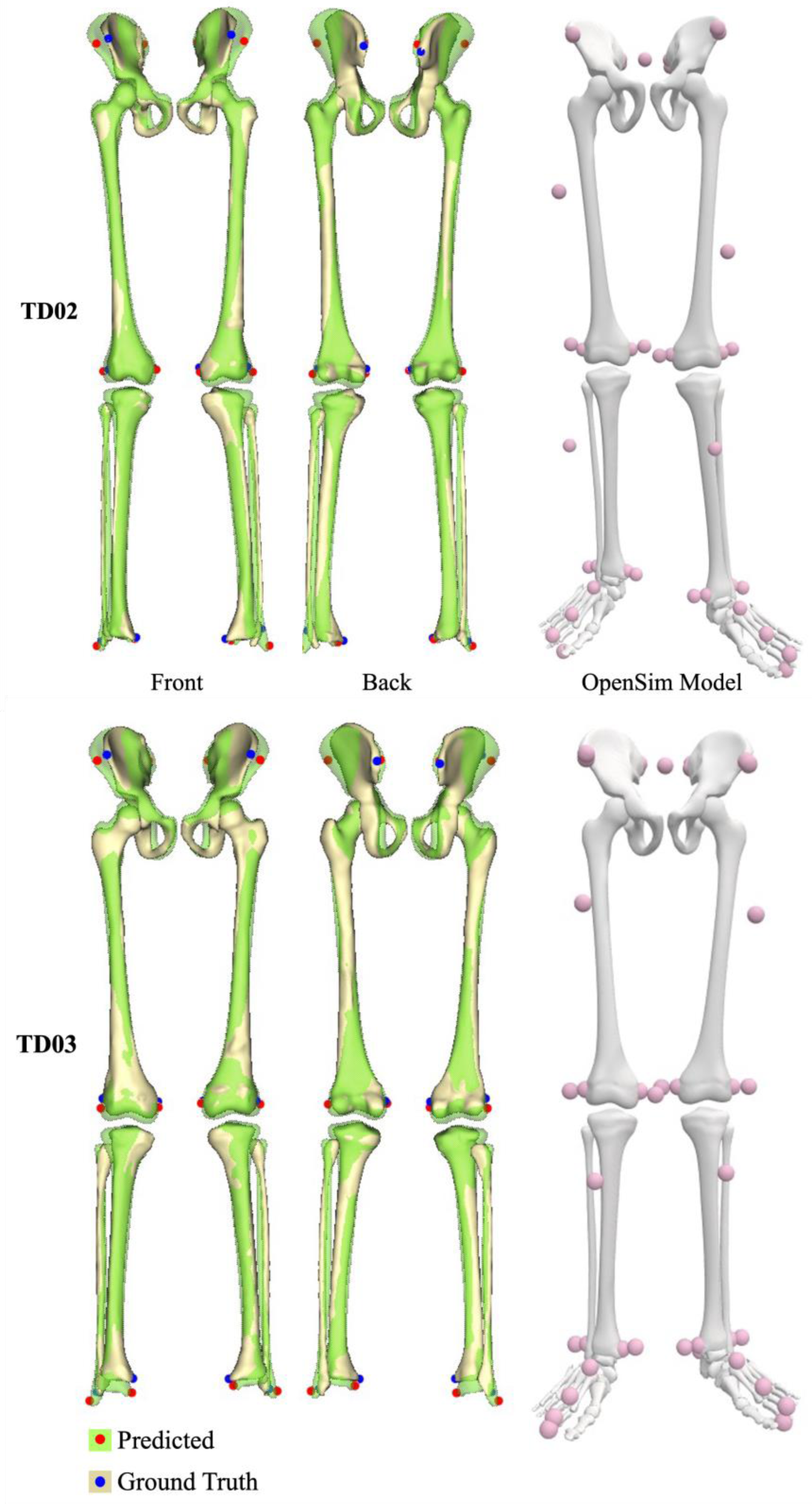
Predicted (green) and ground truth (light yellow) bone geometries for the two test cases (TD02 and TD03) predicted using both-limb-ASM. The generated OpenSim model for each case is shown on the right.

**Table 8:** bone surface error (mm) for prediction of the two example cases (TD02 and TD03) shown as Mean Absolute Error (MAE) ± one standard deviation using the articulated shape model (both-limb-ASM)

|  | **MAE (mm) ± SD (mm)** |  |  |  |  |
|---|---|---|---|---|---|
| **Participant** | **Pelvis** | **Left Femur** | **Right Femur** | **Left Tibia/Fibula** | **Right Tibia/Fibula** |
| **TD02** | 3.41 ± 3.1 | 1.88 ± 1.6 | 1.77 ± 1.3 | 2.51 ± 1.9 | 1.96 ± 1.7 |
| **TD03** | 3.28 ± 2.6 | 1.64 ± 1.2 | 2.07 ± 1.8 | 1.78 ± 1.5 | 1.80 ± 1.7 |

The landmark errors are shown in Table 9 and are also visible in Figure 6. Again highlighting the difficulty of predicting the pelvis geometry with errors ranging from 12.6 to 19.0mm for the ASIS markers and 7.8 to 11.4mm for the PSIS markers. Additionally, high errors were seen in the ankle markers (medial and lateral malleoli) likely indicating discrepancies in marker placement or field of view from the MRI scans.

**Table 9:**
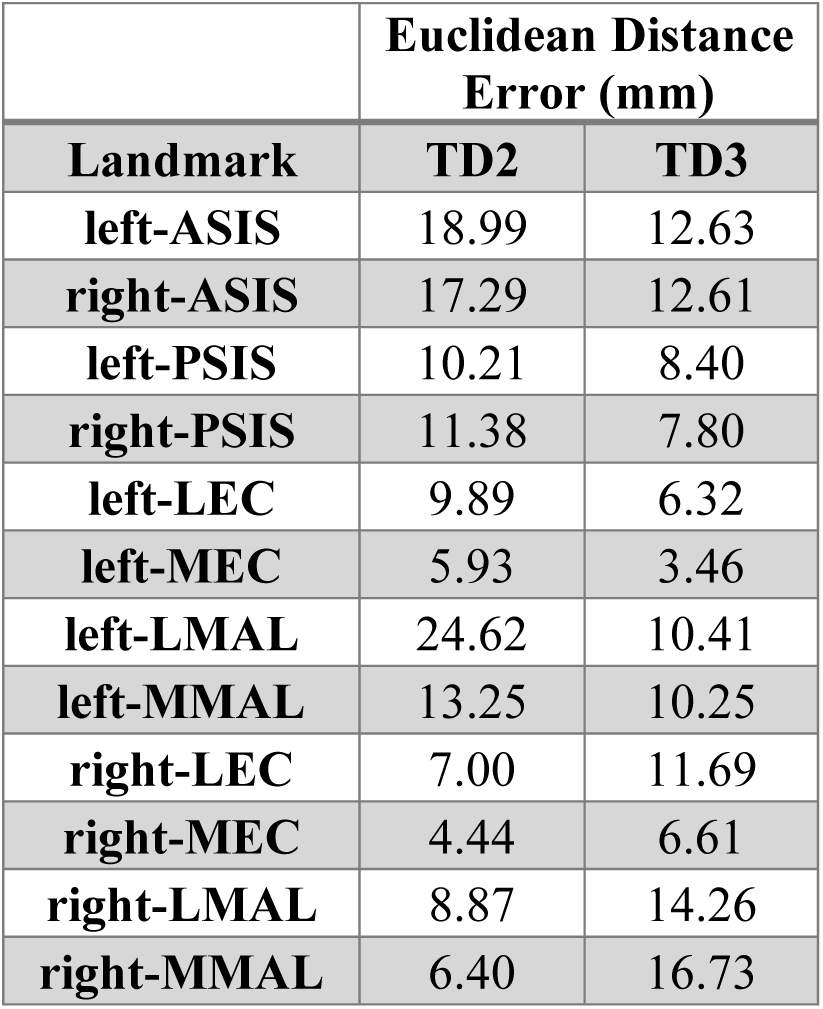
landmark errors (mm, as euclidean distance) for prediction of the two example cases (PLB02 and PLB03) using the articulated shape model (both-limb-ASM). The landmarks include left/right Anterior Superior Iliac Spine (ASIS), Posterior Superior Iliac Spine (PSIS), Lateral Epicondyle (LEC), Medial Epicondyle (MEC), Lateral Malleolus (LMAL), and Medial Malleolus (MMAL).

|  | **Euclidean Distance Error (mm)** |  |
|---|---|---|
| **Landmark** | **TD2** | **TD3** |
| **left-ASIS** | 18.99 | 12.63 |
| **right-ASIS** | 17.29 | 12.61 |
| **left-PSIS** | 10.21 | 8.40 |
| **right-PSIS** | 11.38 | 7.80 |
| **left-LEC** | 9.89 | 6.32 |
| **left-MEC** | 5.93 | 3.46 |
| **left-LMAL** | 24.62 | 10.41 |
| **left-MMAL** | 13.25 | 10.25 |
| **right-LEC** | 7.00 | 11.69 |
| **right-MEC** | 4.44 | 6.61 |
| **right-LMAL** | 8.87 | 14.26 |
| **right-MMAL** | 6.40 | 16.73 |

The automatically generated OpenSim models for each case are shown in Figure 6 showing the ability of the methods to rapidly generate custom OpenSim models.

## 4 Discussion

We have presented a novel workflow for automatically generating paediatric OpenSim models using an articulated shape model of the lower limb. We have expanded our previous model to both the left and right side of the lower limb and evaluated the model’s accuracy when predicting from skin surface landmarks. We showed that our shape model can predict the bone geometry of the pelvis, femur, and tibia/fibula with root mean squared errors of 2.64 ± 0.99mm, 1.91 ± 0.59mm, and 1.69 ± 0.55mm, respectively, in children aged 4 to 18 years. Skin padding and clinical bone measurements were predicted to a reasonable accuracy apart from femoral anteversion angle. We also showed the ability of our workflow to automatically generate a musculoskeletal model from motion capture data.

We developed a random forest algorithm to predict skin padding at 12 common motion capture landmark sites. The lowest RMSEs were observed when predicting from age, height, mass, ASIS width, and left epicondylar width for the left side and age, height, mass, ASIS width, right epicondylar width, and right malleolar width for the right side. This resulted in an R^2^ values of 0.59 and 0.56 and average errors of 2.97mm ± 3.68mm and 2.96mm ± 3.57mm for the left and right side, respectively. This shows that a variety of input data is required to predict skin padding in a paediatric population, beyond just body mass. More accurate prediction results may be achieved with additional input factors, such as body fat percentage, level of activity, and/or pubertal stage. Regional skin padding varied within our dataset with the largest values seen in the pelvis ASIS and PSIS markers ranged from 0.06 to 70.8mm and 5.9 to 87.6mm, respectively, which is where stomach fat tends to accumulate and was much larger than expected, indicating the need for identifying/predicting subject specific skin padding in musculoskeletal simulations. The skin padding values at the femoral epicondyles and medial-lateral malleoli were smaller by comparison with values ranging from 3.9 to 46.3mm at the femoral epicondyles and 0.4 to 17.7mm at the malleoli. These ranges indicate the importance of obtaining patient-specific skin padding values. The large range of values at each landmark add to the difficulty in predicting skin padding in a paediatric population. None of our predictive factors had strong correlations with skin padding, indicating that other factors may be at play. However, using the articulated shape model, bone geometry was still predicted with low errors, highlighting the robustness of the model for predicting bone geometry from skin surface landmarks in a paediatric population. Although there is limited literature on skin padding in children, there are two studies we can draw comparison to. Town et al. found an average skin padding of 2.47 ± 1.2mm and 2.14 ± 1.22mm at the right and left iliac crest of the pelvis (ASIS), respectively, in 100 healthy individuals aged 16 to 35 years [28]. These values are much smaller than our observed average ASIS skin padding of 11.0mm, however, soft tissue skin thickness was measured laterally rather than anteriorly, which may produce some differences. Additionally, a majority of their ‘younger’ dataset (16 to 35 years) was obtained from a relatively fit kinesiology student population. Town et al. also produced regression equations for predicting soft tissue thickness at the pelvis ASIS markers and found an RMSE of 8.6mm, and R^2^ value of 0.704 to 0.718 when predicting from height, abdomen skinfold, anterior mid-thigh skinfold, and upper thigh circumference. While our study achieved much lower prediction errors than this, our R^2^ values were smaller. Derraik et al. found an average combined dermis and subcutis thickness of 11.02mm and 14.89mm in the abdomen and 9.28mm and 14.96mm in the thigh, for males and females, respectively in children aged 6 to 19 years with type 1 diabetes [24]. Notably, these are not at anatomical landmarks, however are similar to the values observed in our study.

When looking at predicted bone surface, we achieved similar results as our previous iterations of the model [22], even though we now predict from skin surface landmarks rather than bone surface landmarks. RMSEs from the leave-one-out analysis of the articulated shape model were 2.64 ± 0.99mm, 1.90 ± 0.58mm, 1.91 ± 0.59mm, 1.69 ± 0.56mm, and 1.68 ± 0.54mm for the pelvis, left femur, right femur, left tibia/fibula, and right tibia/fibula, respectively. Bounding the model and predicting skin padding have enabled these low prediction errors. Our model outperformed the adult articulated shape model developed by Zhang et al. [21] who found bone surface errors of 4.23 ± 1.05mm, 4.57 ± 1.75mm, and 3.63 ± 0.79mm for the pelvis, femur, and tibia/fibula respectively. When comparing to articulated shape models which predict bone geometry from medical imaging data our model performed similarly. Mean errors of 1.1 ± 0.2mm and 1.9 ± 0.2mm for the femur and pelvis were found when using x-ray data [43]. Another model found RMSEs of 1.52 ± 0.44mm and 2.65 ± 1.22mm when predicting the femur and tibia/fibula from MRI data [44]. The articulated shape model still produced far smaller errors compared to linear scaling of the adult OpenSim gait2392 model highlighting the importance of generating participant specific geometries for musculoskeletal models in a paediatric population.

Bony landmark errors predicted from the articulated shape model were quantified as the Euclidean distance between the landmarks on the bone geometry predicted by the articulated shape model and the ground truth landmarks from the CT scan segmented bones. These errors were similar to those seen in our previous articulated shape model [22], with on average a 0.7mm increase in ASIS and PSIS landmark errors, a 0.59mm decrease in lateral epicondyle landmark errors, a 0.71mm decrease in medial epicondyle landmark errors, a 1.14mm increase in lateral malleolus landmark errors and a 0.11mm decrease in medial malleolus landmark errors. This slight increase is likely due to the prediction of skin padding in the model in which we used an RF algorithm. The error in this prediction likely translated through to the bony landmark errors. Additionally, hip joint centre errors were found to be on average 3.76mm, which is slightly higher on average (by 0.81mm), but comparable to our previously developed single bone shape models, where hip joint centre location was predicted using a statistical shape model from demographic factors and linear bone measurements [45]. Overall, the updated model predicted bone surface landmarks well, considering we are predicting from skin surface landmarks.

Bone measurements taken from the predicted bone geometry were reasonably well predicted by our articulated shape model. Larger errors were seen in the femoral anteversion angle (7.43°) and linear bone lengths (5.86mm for the femur and 5.37mm for the tibia/fibula) compared to the other clinical bone measurements. However, our updated model improved on tibial torsion prediction compared to our previous model (by 2.85°). Higher errors in anteversion angle have been found throughout all our previous models, indicating the difficulty of predicting this value without extra input at the proximal femur. Accurate representation of torsional/angular values is important as they influence musculoskeletal model outputs such as joint contact, and muscle forces [7]. This could be remedied by inclusion of data for these regions such as additional landmarks (greater trochanter or tibial condyles) or partial medical imaging data of the hip and knee joint, which can provide extra information for the shape model.

The paediatric test cases illustrated the ability of the workflow to rapidly and accurately generate custom OpenSim models for use in motion capture analysis. The model had some difficulty in predicting the pelvis geometry, specifically the pelvis width. This is likely due to a combination of marker placement error, and the prediction of the skin padding at the pelvis. As we have predicted bone surface markers from the motion capture markers, there is some error in this methodology based on the predictive factors we have access to. The pelvis markers had a large range of values, making the prediction challenging. This could be remedied by including skin surface data as part of the shape model, including partial medical imaging input to give a better indication of bony landmark location, or introducing calliper measurements at input predictors. Also observable in the tibia/fibula were higher errors at the distal end. In the case of TD02, there was some warping of the MRI imaging at the end of its range which could explain this. Overall, we outlined methods for the automatic generation of a paediatric musculoskeletal model from motion capture landmarks by using our articulated shape model prediction to create an OpenSim model.

Limitations of this study include the fact that the landmarks used for prediction were taken from the skin surface of the participant lying in a supine position, rather than an upright position, resulting in skin compression on the posterior side. The imaging dataset to create our shape model was obtained from an urban Australian population; therefore, may not accurately represent a global or specific indigenous population. Additionally, some age groups had fewer CT scans available (e.g., 7 to 10 years old), meaning these age groups may not be represented as well as other age groups in the cohort. Creation of lower limb musculoskeletal models also requires the addition of the patellae and feet, which will be included in future work. One drawback of the articulated shape model was the inability to accurately capture femoral anteversion angle and segmental length values in this population. Future work will involve investigating inputs that can help the model more accurately predict these values, such as partial medical imaging data, additional landmarks, or a physical examination. Finally, the shape model and Opensim model creation tool created for this study is available on the open-source platform SimTK.org (https://simtk.org/projects/paed_ssm) or alternatively on GitHub (https://github.com/musculoskeletal/articulated-ssm, https://github.com/musculoskeletal/opensim-model-creator).

## 6 Conclusions

We have demonstrated the accuracy of using an articulated shape model to predict bone geometry in a paediatric population based on 12 skin surface landmarks. This model outperformed single bone shape models and linear scaling and performed similarly to our previous articulated shape model which predicted bone geometries from bone surface landmarks. Using this articulated shape model, we automatically generated paediatric musculoskeletal models for use in clinical gait analysis. Our workflow will save time in the development of musculoskeletal models without sacrificing the accuracy that is lost when using linear scaling. We have denoted this workflow: Mini Musculoskeletal Automated Pipeline or ‘MiniMAP’.

## Data Availability Statement

Shape models, clinical bone measurements, and skin padding values are available at: https://simtk.org/projects/paed_ssm

Code repositories for generating the articulated shape model and OpenSim models are available at: https://github.com/musculoskeletal/articulated-ssm, https://github.com/musculoskeletal/opensim-model-creator

## Acknowledgments

We would like to thank the Victorian Institute of Forensic Medicine (VIFM), Melbourne, Australia for their support in providing this unique dataset. We would also like to thank the financial support of the University of Auckland doctoral scholarship (New Zealand), the Friedlander Foundation (New Zealand), the Health Research Council of New Zealand, the Aotearoa Foundation (New Zealand), and the Medical Research Future Fund (Australia).

## Author contributions

L.C. performed the code development, data processing, and analysis of the data and wrote the first draft of the manuscript. J.C. created the study and obtained the data. J.C. and T.F.B. assisted in creating and reviewing the manuscript. T.S.W assisted in code development.

## Conflict of Interest Statement

The authors declare no competing interests.

## Notes

### Competing Interest Statement

The authors have declared no competing interest.

